# Application of subject-specific adaptive mechanical loading for bone healing in a mouse tail vertebral defect

**DOI:** 10.1101/2020.09.13.295402

**Authors:** Angad Malhotra, Matthias Walle, Graeme R. Paul, Gisela A. Kuhn, Ralph Müller

**Affiliations:** Institute for Biomechanics, ETH Zurich, Zurich, Switzerland

## Abstract

Methods to repair bone defects arising from trauma, resection, or disease, continue to be sought after. Cyclic mechanical loading is well established to influence bone (re)modelling activity, in which bone formation and resorption are correlated to micro-scale strain. Based on this, the application of mechanical stimulation across a bone defect could improve healing. However, if ignoring the mechanical integrity of defected bone, loading regimes have a high potential to either cause damage or be ineffective. This study explores real-time finite element (rtFE) methods that use three-dimensional structural analyses from micro-computed tomography images to estimate effective peak cyclic loads in a subject-specific and time-dependent manner. It demonstrates the concept in a cyclically loaded mouse caudal vertebral bone defect model. Using rtFE analysis combined with adaptive mechanical loading, mouse bone healing was significantly improved over non-loaded controls, with no incidence of vertebral fractures. Such rtFE-driven adaptive loading regimes demonstrated here could be relevant to clinical bone defect healing scenarios, where mechanical loading can become patient-specific and more efficacious. This is achieved by accounting for initial bone defect conditions and spatio-temporal healing, both being factors that are always unique to the patient.

## Introduction

The management of critical-size bone defects continues to present surgical challenges. Trauma and bone resection can lead to lengthy recovery times or amputation. The use of autografts is the current gold standard, however, is quantity-limited and accounts for 20% of the complications^1^. Despite advances in biomaterial development and understanding of signaling mechanisms, the search for improved treatment methods of such bone defects continues.

One treatment method of interest is mechanical loading of bone. Mechanical interactions have a long-established relationship to bone physiology, leading to earlier concepts of micromotion during bone defect healing^2^, and the significance of fracture instability in healing outcomes^3,4^. The effects of mechanical forces on bone healing have been previously reviewed^5,6^; mechanical loading is likely to depend on frequency^7^ and cycle number^8^, influences mesenchymal stem cell differentiation^9^, and has a role in guiding healing towards primary and secondary bone healing pathways^10^. In contrast, mechanical loading and relative motion of fragments showed little to no benefit in other studies^11,12^, though the timing of the changes of the mechanical environment in relation to healing phases could also play a role^13^. This further highlights the need to understand the mechanical environment in and around bone defects during healing. This mechanical environment of bone includes the relevant surrounding hard and soft tissues, and their interaction as subject to Newtonian mechanics, which allows computational exploitation for assessing the loading history and relationship to morphological changes^14^. However, the translation of these load-driven bone (re)modelling concepts to highly unique bone defect healing scenarios is lacking. Another current challenge that arises is how to determine the force that needs to be applied in a subject-specific manner in order to have a maximal mechanobiological cue without damaging the bone.

Finite element (FE) analysis is a well-proven approach to understand micro-scale strains and has been previously successfully used to correlate strain and in vivo bone (re)modelling activities in mice^15–17^. In such workflows, bone mechanoregulation can be studied non-invasively by combining imaging and computational FE-derived strain estimation methods^18^. With advances in computational power, the time needed to calculate voxel-based strains relevant for bone healing has drastically shortened in the last decade. Depending on the complexity and resolution deemed suitable, it is possible to do this immediately after imaging to limit unnecessary or additional handling and anesthesia in mice. This concept is concurrently presented in a mouse femoral defect model^19,20^, and in combination introduce the concept of real-time finite element (rtFE) analysis to describe this approach. Since mechanical loading is effective only within a certain strain window^21^, maintaining the applied loading within this window is critical. Loading too high risks damage (Supplementary Fig. S1a) and loading too low risks no effect. FE analysis can be used to estimate the optimal forces, and the rtFE method builds on this by streamlining the process of imaging, analysis, and treatment.

Two factors are relevant for determination of the appropriate loading conditions: the defect itself; and its changes due to healing. First, every defect is unique in shape, size, and location. This, in combination with the surrounding structural anatomy, will influence how strain is transferred across the region. Second, inherent differences between an individual’s healing capabilities will always exist. When bone heals, the tissue-level stiffness increases. If constant stimuli are applied, micro-level strains will decrease as bone forms and reinforces the defect. This would lead to a lowering of the regional strain, and result in sub-optimal loading conditions. Therefore, applying individualized and adapting loading regimes that factor in the defect and its unique healing has high potential to promote bone healing, not only in mice, but in humans, where these two factors can vary widely case to case.

Applying loads to a bone defect, and imaging it at the necessary resolution, are fundamental requirements of this study. A previously described method enabled the loading of mouse vertebrae^22–24^, and importantly, allowed high-resolution scanning of the vertebrae. This was recently advanced on to investigate the loading frequency effect on mouse vertebral bone parameters^25^. To investigate specifically bone defect healing in this current study, a caudal vertebral bone defect model was developed, which could be incorporated into the previously successfully used workflows. This model is straightforward, reproducible and facilitates biomaterial placement^26^. In comparison to commonly used mouse hind leg models, the vertebral model can reduce the complexity and uncertainty within the FE simulation. For example, hind leg models can have more complex boundary conditions due to more complex joint constraints of the limb, greater influence from external bodyweight loadings due to gravity, and higher internal muscle-bone loads applied by the mouse during ambulation^27^. Furthermore, the vertebral model requires no surgical fixation, and allows for early loading post-surgery due to the high initial stability of the defect region compared to osteotomy-based models. This lack or presence of fixation can make comparisons difficult, even if one considers that the mechanobiological responses are consistent. Overall, the vertebral defect model introduced here provides another anatomical location to advance theories of bone mechanoregulation during defect healing.

Longitudinal time-lapsed imaging with micro-computed tomography (micro-CT) is a suitable method to image bone at the early stages of healing^28^. This can provide short time interval snapshots of the healing progression and create high-resolution datasets for FE simulations. This time-lapsed imaging method is currently most feasible in smaller animals, and has been previously demonstrated in vertebrae^22^, and recently within both mouse femoral defect models^29^, and to assess longer-term morphological changes of intact mouse vertebrae^25,30^. This method is highly relevant here to provide weekly insights into the subject-specific healing progression, and when used in combination with the rtFE methods, it enables in vivo assessment and proportional changes to the loading conditions simultaneously.

The objective of this study was to test the feasibility and effect of subject-specific adaptive mechanical loading to treat bone defects. This was investigated in a newly developed mouse vertebral defect model that allowed mechanical loading across the vertebrae, as well as high-resolution imaging for rtFE analyses. As humans also share similarities to bone loading responses^31^, such a workflow could pave the way for patient-specific loading regimes that increase the effect and repeatability of mechanical loading regimes for bone regeneration strategies.

## Results

### General observations

All mice tolerated the defect surgery procedure well, with no impaired movement or indications of pain, except one, which was euthanized out of precaution immediately after surgery due to perceived excessive bleeding during surgery. None of the remaining mice experienced adverse effects from the rtFE procedure, with no fractures or additional pain due to the workflow. One additional mouse was euthanized after one week due to increasing and persistent swelling and signs of osteolysis in the adjacent vertebrae around the pins. Furthermore, one defect was excluded from the data analysis because it was drilled through two cortices. In total, this resulted in groups of 6 (control) and 7 (rtFE loading) mice.

The rtFE method implemented in the study increased the total anesthesia time from approximately 22 minutes for the classic procedure to approximately 30 minutes. This additional time was due to the computing time and adjustment of the loading device, and is a downside of the rtFE method (Supplementary Fig. S3). No difference in the time to regain consciousness was noted, and all mice recovered from the anesthesia as expected. Force increased significantly over time (F(3,18)=25.8, p<0.001), with the initial average peak-to-peak force of the cyclic load calculated for the rtFE loading groups was 4.3N (±0.7), and significantly increased per week, first to 4.5 N (±0.5), then to 4.8 N (±0.4), and to 5.2N (±0.3) in the final week of loading.

Healing progressed appositionally from the ventral and lateral internal surfaces of the bone (Fig. 1a). There were no signs of cortical bridging. Dense trabecular bone formed in regions where it would bridge to adjacent surfaces (Fig. 1b), stabilizing the defect against the applied loading and consequent bending moment induced by the defect asymmetry.

**Figure 1.**
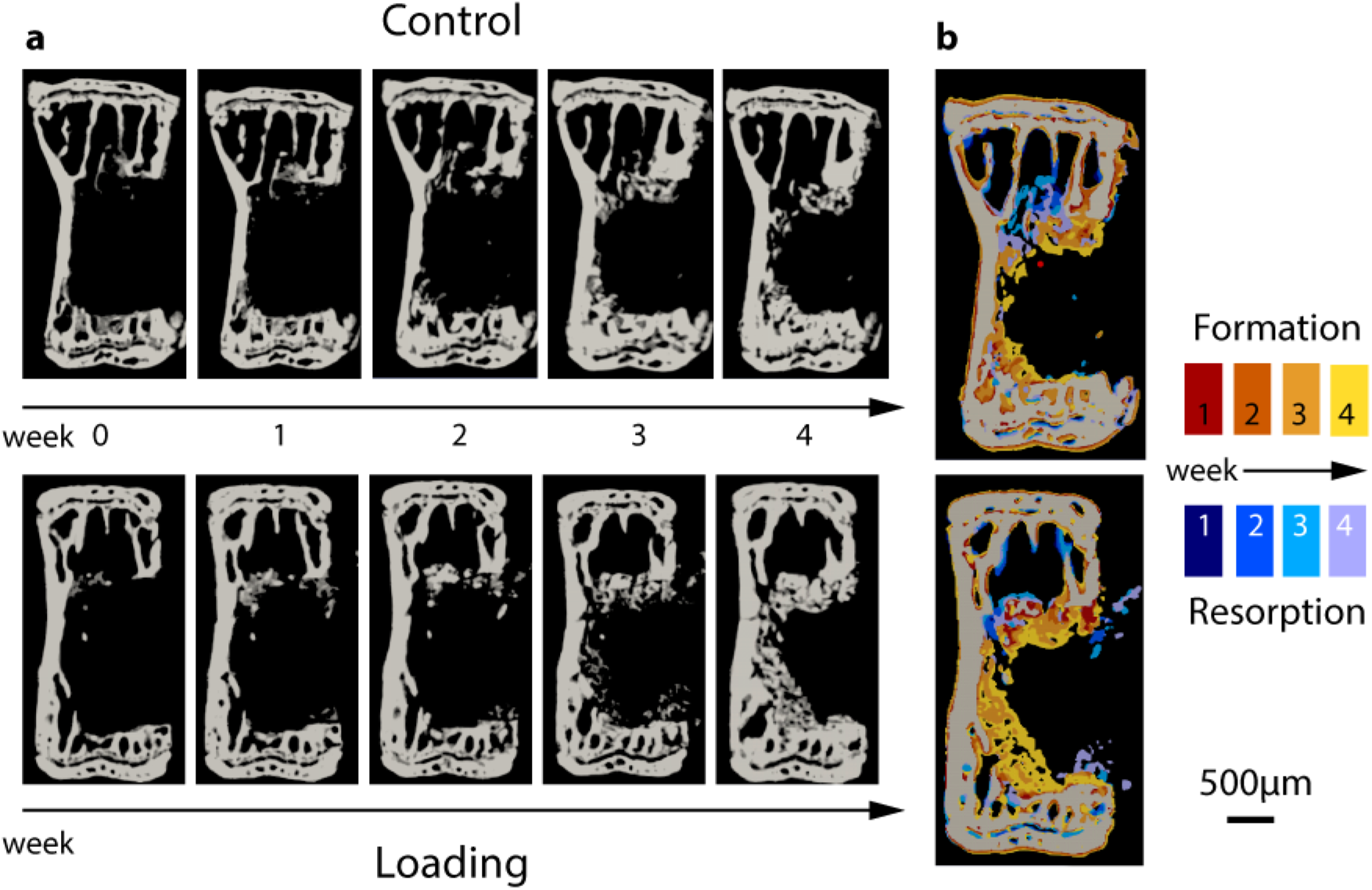
Time lapsed imaging and overlay of formation and resorption on a weekly basis. **(a)** Time lapsed images of representative animals from both groups. **(b)** Weekly overlays show formation and resorption patterns from 1 to 4 weeks post-surgery. Bone formed within the defect without cortical bridging. Red/yellow: bone volume formed at week 1, 2, 3, and 4. Blue/purple: bone volume resorbed at week 1, 2, 3, and 4.

### Longitudinal assessment of bone defect healing

Two volumes were created to differentiate between the initial empty defect space, and the existing surrounding bone. These were the defect centre (DC) and the defect periphery (DP). (Fig. 2a). Within DC (Fig. 2b), a significant interaction was present for bone volume fraction (BV/TV) between time and loading (F(1,4)=8.90, p<0.005)). At week 2, 3 and 4, loading significantly increased BV/TV over controls (p<0.05). BV/TV increased significantly over time in both loading and control groups (p<0.005). In the DP, both time (F(1,4)=164.7, p<0.005)) and the loading (F(1,4)=5.34, p=0.041) had significant overall effects on BV/TV (Fig. 2c).

**Figure 2.**
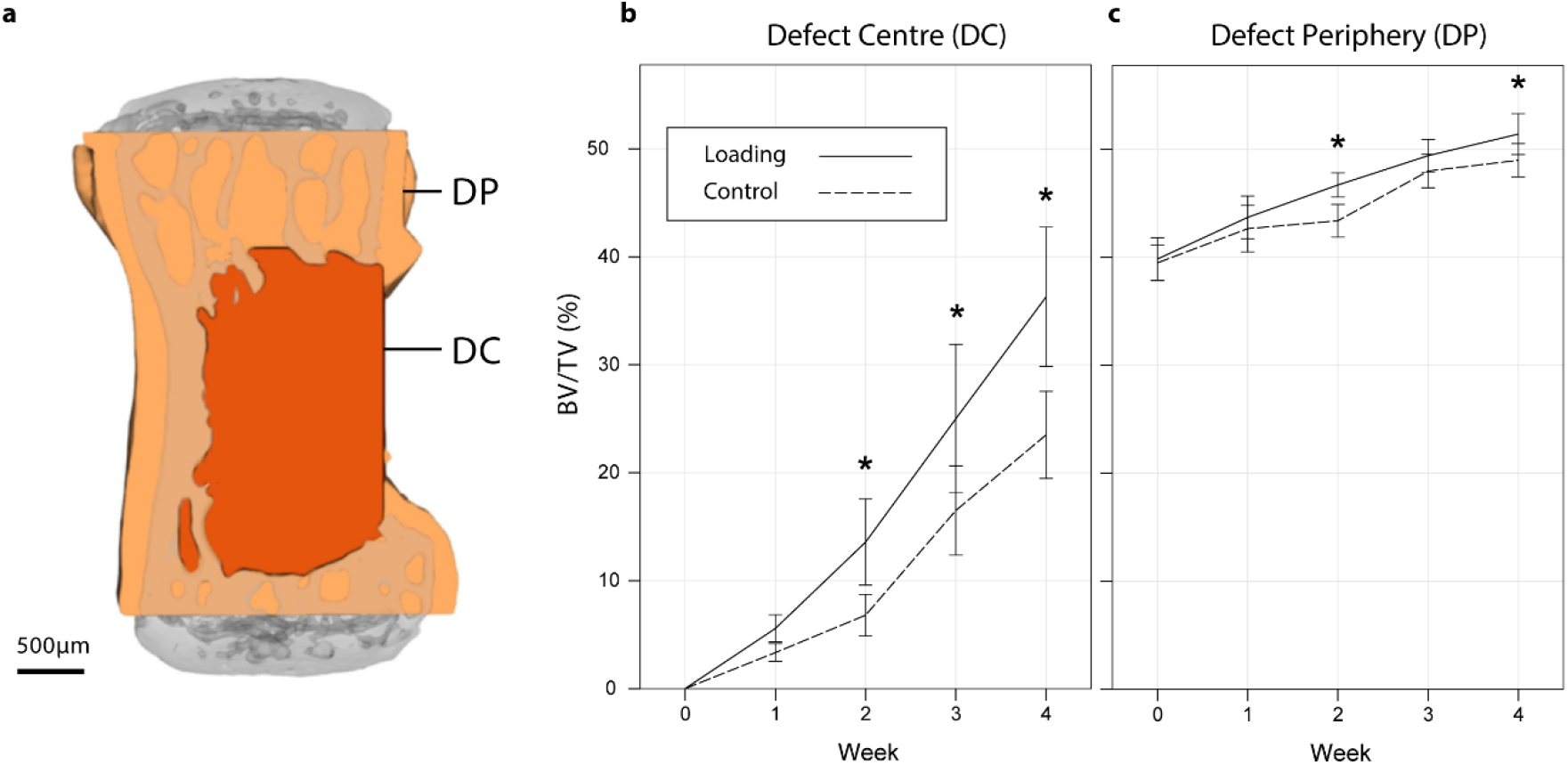
Longitudinal changes in bone volume fraction. **(a)** The vertebrae were divided into a defect centre (DC) and defect periphery (DP). **(b)** BV/TV within the DCsignificantly increased with the rtFE loading from week 2 compared to controls. **(c)** BV/TV within DP was also found to be influenced, but not to the same magnitude or extent as the DC.

Overall, loading (F(1,3)=12.6, p<0.001)) and time (F(1,3)=23.2, p<0.001) had a significant effect on the DC bone formation rate (BFR/DC). Loading significantly increased BFR/DC over controls between weeks 1-2 (F(1,3)=5.83, p=0.013), and weeks 3-4 (F(1,3)=4.39, p=0.042), though did not reach significance for weeks 2-3 (F(1,3)=2.05, p=0.159) (Fig. 3a). Loading did not largely affect the DP bone formation rate (BFR/DP) or DP bone resorption rate (BRR/DP) at any time (Fig. 3b). Also, overall, loading did not have a large effect on BRR/DC or BRR/DP compared to control mice (Fig. 3).

**Figure 3.**
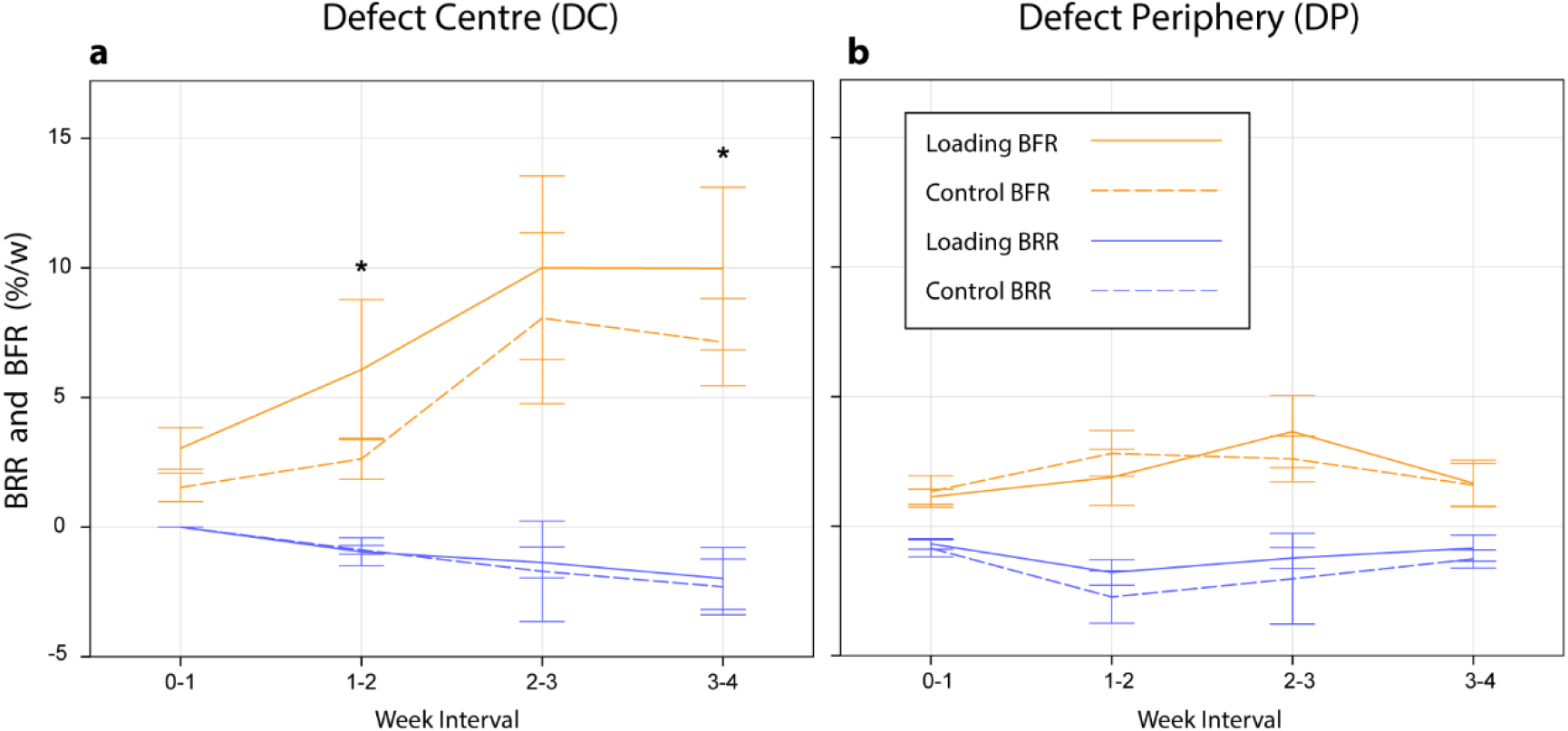
Longitudinal changes in formation and resorption volume fractions. **(a)** Loading influenced the BFR/DC compared to controls, and reached significance at postoperative weeks 2 and 4, while loading did not appear to influence BRR/DC compared to controls. **(b)** Loading did not significantly influence either BFR/DP or BRR/DP compared to controls, at any time interval.

### Mechanics of bone volume changes

After 4 weeks, none of the vertebrae had recovered their pre-surgery axial stiffness based on the applied forces. From week 2 onwards, the treatment had a significant positive effect on FE-calculated stiffness (p<0.05, Supplementary Fig. S2a). There was also a significant positive Pearson’s correlation between BV/TV in the DC, and FE-calculated normalized stiffness (r=0.907, n=51, p<0.005, Supplementary Fig. S2b), while in the DP the correlation to normalized stiffness was significant, but lower (r=0.809, n=51, p<0.005). A pattern was noted in the probability of formation, quiescence, or resorption events within the combined DC and DP regions; they were largely related to effective strain (EFF). The effective strain as a percentile of the max effective strain (EFF/EFF_MAX_) was used to find the conditional probability of a (re)modelling event occurring due to strain. During the first week of healing, bone formed with a random conditional probability (cp(x) = 33.3%). However, in subsequent weeks, bone formed in the upper half of the strain field (cpF(x > 50 %) > 33.33 %). The probability for resorption was the highest within the first 30% of occurred strain leading to a small strain window where bone was predominantly quiescent between 30% and 50% (Fig. 4a). This pattern also occurred in control animals (Fig. 4b) where physiological strains were simulated with FE analysis (Fig. 5). When comparing the loading and control cases, the curvature of the formation, resorption and quiescence profiles in the loading cases had steeper and more pronounced curve sections compared to the control cases. There was also a small shift towards lower EFF/EFF_MAX_ being formative with loading.

**Figure 4.**
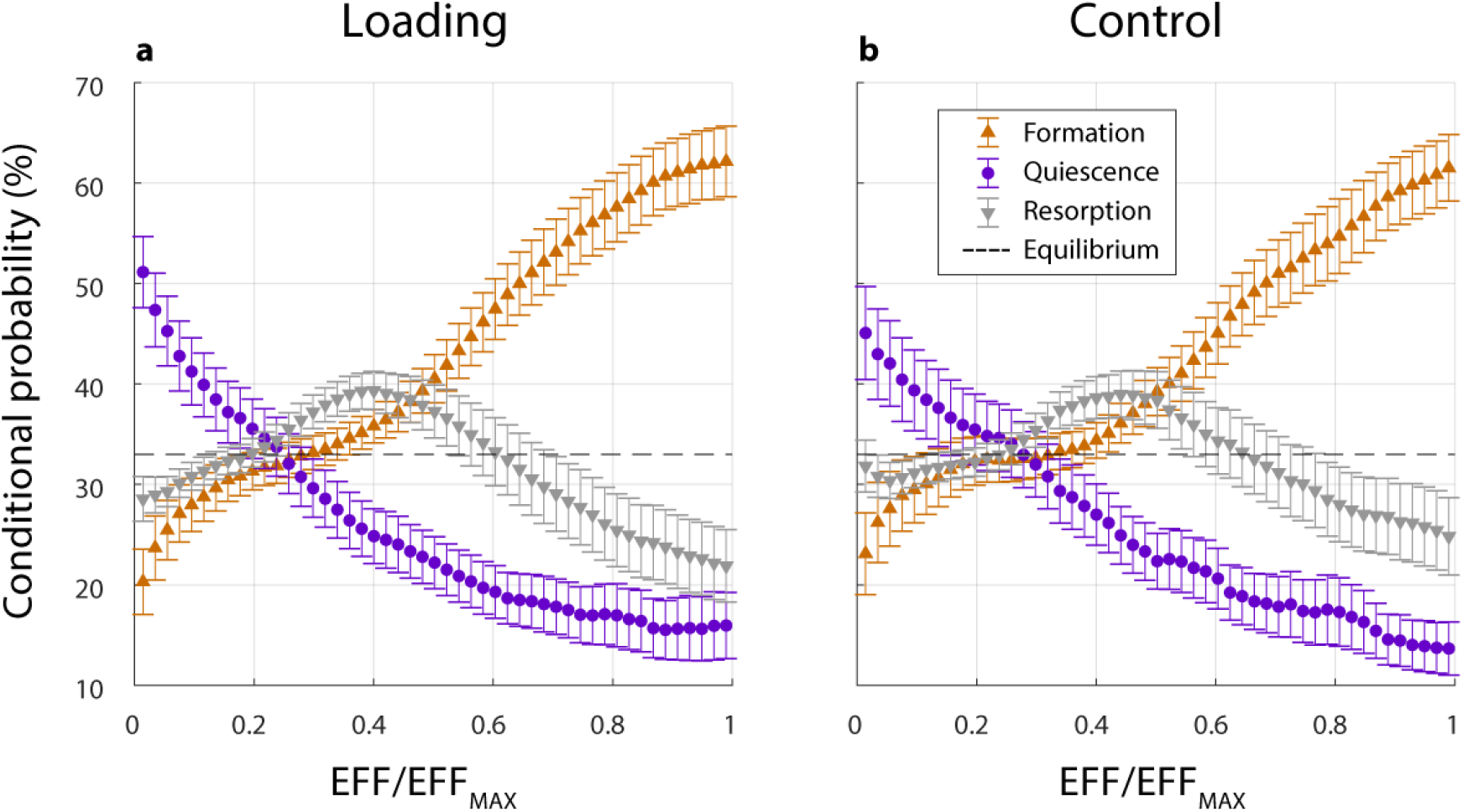
Conditional probabilities of formation and resorption events within the combined DC and DP regions. Effective strain as a ratio of the maximum effective strain value (EFF/EFF_MAX_)). **(a)** Higher ratios of EFF/EFF_MAX_ led to formation activity in treatment groups, and lower ratios led to resorption activity, regardless whether externally loaded in the treatment group, or in **(b)** control animals with an assumed axial strain.

**Figure 5.**
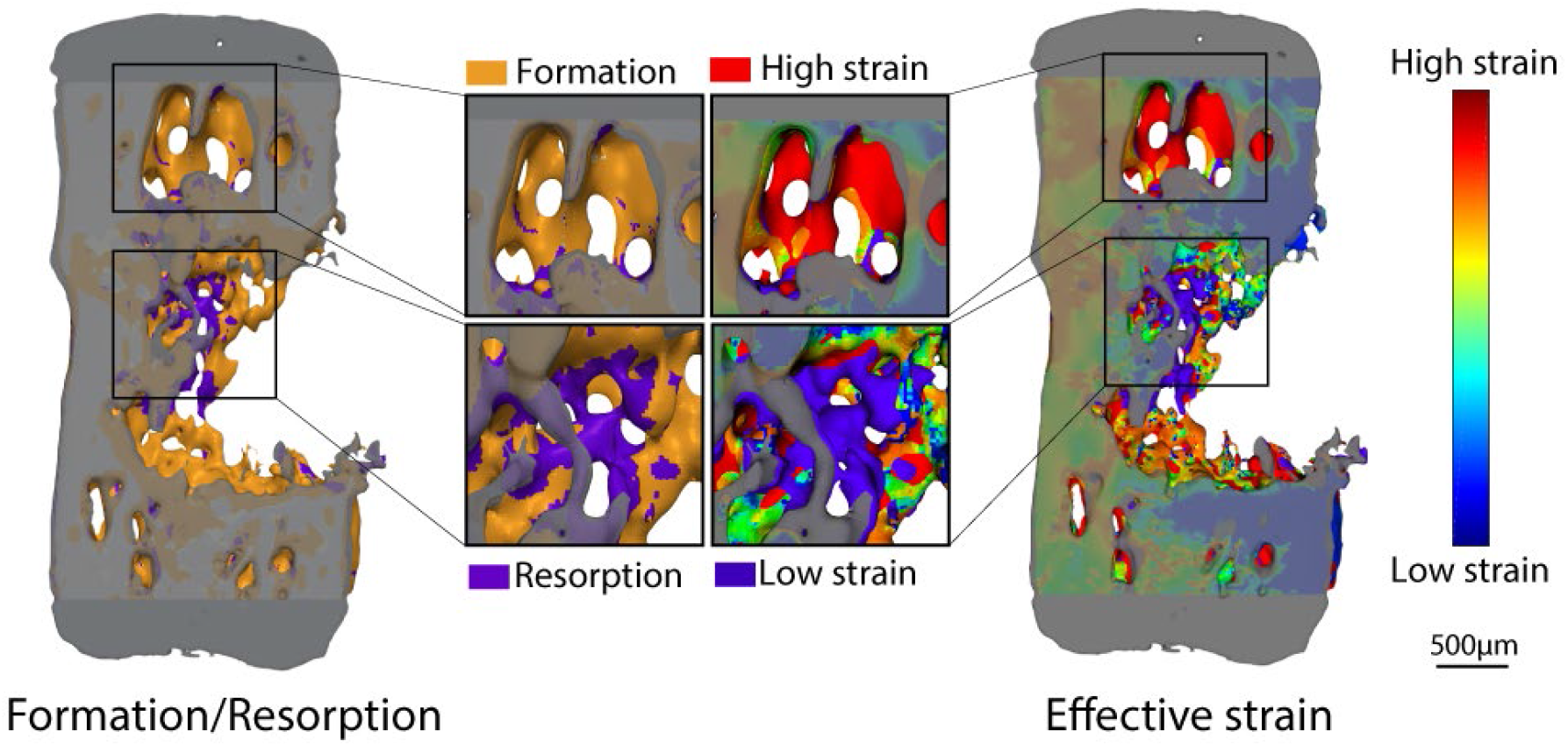
Representative sample depicting formation and resorption relationship with effective strain. Regions of higher effective strain tended to formation, while lower effective strain tended to resorption.

## Discussion

While the influence of mechanical loading on bone (re)modelling is known, implementing this to defect healing creates other challenges. These arise as two aspects, being the changed mechanical environment due to the initial defect, and its unknown healing thereafter.

This study showed that by introducing an rtFE approach to an existing loading set-up^22^, bone defect healing could be significantly improved over no-treatment controls. Importantly, this approach succeeded in avoiding any incidence of fracture due to overloading, and in principle, homogenized strain across different defect shapes, sizes, and healing progressions. Adaption of the loading within the mechanical environment is not novel in itself, with mixed reports on the effectiveness of dynamization^12,32^, in which a change in stiffness of external fixators is adapted over healing periods. Adding to these existing concepts, this study estimated individualized loads to apply during healing; the rtFE approach allows much greater accuracy in the control of strain, as opposed to generic or pre-determined adaptive regimes.

Loading started at two days after the defect was created, at which time the mice were still relatively young at fourteen weeks old. The bone response to loading has previously been shown to have some age-dependency on mice around this age, where six week old mice had a more exaggerated response to loading compared to ten and sixteen week old mice^33^. Hence, it could be questioned whether the positive effect found in this study would also be repeated in older mouse. In this regard, it has been reported that after sixteen weeks old, aging has less of an influence on the response to loading^30^. At fourteen weeks old, the mice in this study are near the border of this apparent age threshold. As such, prior studies suggest the positive effect of this rtFE loading could be beneficial into adulthood and beyond, though further studies would be needed to confirm this.

The timing of loading after bone injury is a topic of debate. In general, loading is known to influence cell activity and function due to tissue deformation and fluid flow^34^. Furthermore, it influences both spatial and temporal biological responses at multiple scales^35,36^. This aspect is more important in the case of bone healing, where multiple overlapping phases exist. Early loading from two days onwards has been reported less effective than delayed loading at two weeks^37^. However, it has also been reported that early cyclic loading may increase oxygen transport to the defect region^38^, promoting the longer term regeneration response. As differences in bone parameters were already noted after two weeks in this study, it seems that early loading of three sessions per week is potentially more effective than delayed loading, especially as subject-specific adaptive loading will be in an efficient range without risking damage to early callus structures.

The new bone that formed within the defect did not appear to be simply recreating the pre-defect bone structure, but to be forming based on other factors. During the defect healing, strain appeared to guide bone (re)modelling activities, more so than an inherent sense of prior bone anatomy. Bone appeared to form in compensation for the asymmetry of the defected bone, without cortical bridging, but with densely arranged trabecular bone within the marrow cavity in a conical, V-like shape. This formation pattern supports the idea of a univerisal cellular mechanobiological response regardless of the location within the body, or the presence of fixation implants. This aligns with concepts of bone resorption being caused by either disuse or stress-shielding, where the cells respond to the loads they experience depending on their mechanical state, and not of the cause of the change. Further, the axial mechanical stiffness over the four weeks, as assessed with FE, did not recover to its pre-defect strength, suggesting that healing would continue into the future. The model was newly developed, and the level of impairment over longer time periods would require further characterization of the model. As this study duration was only four weeks, it is unclear whether (re)modelling would eventually result in a structure similar to the native vertebra over time, or even have healed completely given more time. Regardless, bone formation during early healing was related to and guided by bone strain (Fig. 5). This applied to both the loaded and the control group, which could be exposed to physiological strain that may peak at 4N^14^. Strain-induced bone (re)modelling principles have been previously recognized^15,16^, in which loading favors formation over resorption^17^.

This principle is evident elsewhere, where in a study of mouse femoral fractures, fracture site remodeling after three weeks has shown to be consistent with previously considered remodeling theories^39^. However, such comparisons to other locations within the body can be confounded by non-biomechanical factors. For example, the difference in the presence of bone marrow and stem cells, cancellous and cortical bone ratios, and the large muscles surrounding the femur that supply blood and cytokines. While different locations in the body have different factors that can influence healing, similarities do exist between different bone defect models. This study developed and used a confined partial defect model. When comparing this healing to an externally-fixated mouse femoral full osteotomy defect model, mechanical loading from the fourth week onwards also significantly increased BV/TV in the defect centre region. One difference though was the patterns in the BFR and BRR once loading began^19^. Of relevance here is the phase of healing in which loading begins. In the femoral full osteotomy model, a four week loading delay was implemented to allow for bridging across the defect void. Thus, the inflammatory phase of healing has passed when loading begins. In comparison, the partial vertebral defect used in this study allows for loading within the first week after defect creation, during the early inflammatory phase of healing. When comparing healing in the controls in the weeks following defect creation, the BFR and BRR showed similarities in responses between both the osteotomy and partial defect models. It is clear that many confounding factors exist when making such comparisons, including the timing of loading, the healing phase which loading begins, the anatomical location and defect differences. Despite these differences, the bone healing responses to loading within the defect centre and periphery were comparable between these very different defect types. Overall, this further supports a universal relationship between bone healing and loading in mice. Future experiments could confirm this by extending the timeline of loading past four weeks in this vertebral model, by starting loading in later weeks, or by creating a full osteotomy variation of this partial defect vertebral model. Comparisons to intact vertebrae also support the relationship between healing and loading. The conditional probabilities of formation, quiescence and resorption (Fig. 4) in this defect study correlate with prior studies of intact vertebra, where (re)modelling events have a relationship to the strain percentile, and that loading may slightly shift the strain percentile in which formation events occur^15,25^. Considering the above studies, this rtFE study also supports these mechanoregulation theories and further validates the principle in mouse vertebral bone defect healing as well.

Over 80% of the variance (r=0.907, R^2^=0.823) in BV/TV could be accounted for by change in FE-calculated normalized stiffness, and this demonstrates the rtFE approach’s ability to estimate the loading intensity that should be applied. This provided some validation of the rtFE methods used. The FE method, however, only accounted for purely axial compression, aligned to the principal component of the vertebra. In reality, the vertebra was able to vibrate in different modes as it would be constrained differently to the FE models. Therefore, it would have inevitably had some external bending and rotational components not factored in by the simplified FE analyses. Despite this, the relatively high correlation provides suitable confidence in the rtFE protocol. It kept the actual computational processing time to around two minutes, to avoid the well-known effects of animal anesthesia and its possibility to confound the results^40^. Additionally, this correlation is noteworthy considering a dynamic in vivo load was simplified to a static linear simulation. Such linear simulations have previously been reported to be appropriate^41^ and capture this dynamic behavior via a static apparent modulus^42^. Meanwhile, future improvements to computational non-linear analyses may provide future insights^43^. Overall, while assumptions and simplifications exist, this approach was able to balance computational accuracy and cost, and provide confirmation of the usefulness of the rtFE method.

This study did not compare subject-specific adaptive loading group to any traditional, non-adaptive, non-subject-specific loading group. Mechanical loading is well-established to enhance bone healing, and this has been extensively demonstrated in a variety of animal models using various loading modalities. But many questions remain on how to implement this knowledge into practice, where defects and their healing progression can vary widely. This study developed and implemented an objective 3-dimensional imaging and analysis method to assess a defect and its healing, and demonstrated how this could be linked to a known effective loading regime while reducing secondary fracture risks (Supplementary Fig. S1). This overall approach attempts to foresee technological progress and tools that could be more reliable than, for example, subjective grading scales of fractures based on 2-dimensional imaging with subsequent loading based on this assigned grading scale. In this regard, this study does not provide evidence that the complex, objective methodology presented in this study provides improved outcomes compared to a simpler subjective analysis and/or non-adaptive loading regime. Future studies could be designed to investigate if such a benefit truly exists between these approaches. In principle, though, a leaning towards objective, (semi-)quantitative analyses have historically prevailed over subjective, qualitative analyses, and this study attempts to follow this path.

As discussed, assumptions and simplifications created several limitations to this study which cover both animal and computational aspects. For the animal aspects, the mice were relatively young, the healing was not completed within the four weeks, and three mice were excluded which reduced study power. Defects were created using relatively basic tools, and while this provides simplicity and an ability to apply the model, it also introduces some variability in the defect volume across animals. However, this is factored for in the BV/TV calculation. The pinning of the adjacent vertebra also lacked certain control in position and angle, which creates unknowns in how the defect vertebra is loaded, and the modes it vibrates in considering the semi-constrained nature. Future studies could be extended past four weeks in older mice, improve the defect precision and repeatability, and increase pinning control. As for the computational aspects, one of the greatest limitations is the simplifications in the FE model. Firstly, the materials and models are linear elastic, which would not capture any non-linear behaviour of vibration or visco-elastic effects. Material properties of the modelled discs were that of bone, and all surrounding and void voxels not designated as bone were assigned a Young’s modulus of 3MPa, including what would be muscle or air. As such, both the disc and non-bone regions are therefore stiffer than reality. Many of these computational limitations relate to the micro-FE solver, ParOSol^44^; however, these compromises enabled the efficient use of running the FE remotely on a supercomputer, which created the possibility for near real-time results. While computational power is itself a limitation, future studies could explore methods to more accurately model such a dynamic system, and further validate the simplifications and assumptions used.

To translate this approach to patients, not only would further research and development be needed, but for technological advancements also to continue. Most obviously, the computational and hardware technology used in this study is not currently available to clinicians. Secondly, any patient-specific solutions within the FE-realm require computational assumptions to be made, which requires further expertise when applying case-by-case. These current limitations in translatability will likely be overcome as technology develops; this study demonstrates the possibilities the future research can strive towards, once technology and methods inevitably catch up for use in the larger-scales needed for humans.

In conclusion, individualized real-time adaptive loading can be achieved through a combination of micro-CT imaging, followed immediately by FE-solved strain distribution, and finally rescaling and application of a cyclic loading force accordingly. Further investigation is needed to compare this to traditional non-adaptive methods. This rtFE approach is highly relevant for clinical scenarios where bone fractures and their healing progression are unique. This approach optimizes loading intensity, and has the potential to reduce the risk of re-fracture or ineffective mechanical loading, thus improving the healing of bone defects.

## Materials and Methods

### Study design and surgery

Approval was obtained for the animal experiments from the cantonal ethics committee from the Kantonales Veterinäramt Zurich (Zurich, Switzerland, ZH029/18) prior to the study, and all experiments were performed in accordance with Swiss animal welfare act and ordinance, and ARRIVE guidelines. The study included two groups: an rtFE loading group, that were adaptively loaded (3.2-5.5N, 10Hz, 5 minutes, 3000 cycles), and; a control group, that received sham loading (0N) and similar handling. Groups were allocated by block randomization within a larger study, with sample sizes estimated from previous similar research within the laboratory^15^, in which 2 groups for a repeated (4) measures ANOVA using G*Power (β=0.8, α=0.05, f =0.7, number measurements=4, correlation=0.8) estimated a total sample size of 16 (n=8 per group). All surgical, scanning and loading procedures were performed under isoflurane anesthesia (induction 5%, maintenance 1-2%, in O2). To be able to apply loading, three weeks prior to defect surgery, stainless steel pins (Fine Science Tools, Heidelberg, Germany) were inserted in the fifth and seventh caudal vertebrae under fluoroscopic control, as previously described (4). Perioperative analgesia (25 mg/L, Tramal, Gruenenthal GmbH, Aachen, Germany) was delivered via the drinking water for pre-emptive pain relief two days prior to the defect surgery, and for three days post-surgery. All surgeries were performed by the same surgeon. For both groups, defects of approximately 0.8mm x 1.5mm were placed on the dorsal surface of the sixth caudal vertebrae of female fourteen-week old C57Bl/6JRj (Janvier Labs, Saint-Berthevin, France) mice using an electric rotary drill (Micro Drill, Harvard Apparatus, Holliston MA, United States) with 0.6mm and 0.8mm burs. This created an elongated void running along the dorsal aspect of the vertebrae (Supplementary Fig. S3). Humane endpoints included fracture, infection, bodyweight loss of > 15%, or inability to freely eat or drink.

### Imaging and finite element methods

Vertebral defects were scanned at 10.5 μm resolution on the day of surgery, and weekly thereafter, using an in vivo micro-CT (vivaCT 40, Scanco Medical AG, Brüttisellen, Switzerland, 55 kVp, 350 ms integration time, and 145 μA). The resulting images were used as input for the rtFE procedure for animals in the loading group. Loading mice were kept under anesthesia during the image reconstruction and FE calculation.

The reconstructed images were Gaussian filtered (sigma 1.2, support 1) to reduce noise, and thresholded to assign material properties. Voxels within the threshold range from 395 mgHA/cm^3^ to 745 mgHA/cm^3^ were regarded as bone. This bone was assigned isotropic linear elastic material properties with a Young’s modulus between 4 GPa and 12.8 GPa, in threshold steps of 25 mgHA/cm^3^, in proportion with their density ^45^. The Young’s modulus of soft tissue was set to 3 MPa for values lower than 395 mgHA/cm^3^. The Poisson’s ratio was set to 0.3. Vertebra geometry was aligned to the principal axis of the coordinate system in the z-direction. To achieve an even force distribution across the bone and to counter numerical issues with the solver, discs of 1.68 mm diameter with a Young’s modulus of 12.8 GPa were added at the distal and proximal ends of the vertebrae, similar to previously used^17^. The outer surface of the distal disc was fixated using Dirichlet boundary conditions. The outer surface of the proximal disc was deformed by 1% by applying a pure compressive force in the normal (z) direction. Each model was solved using the ParOSol solver^44^, running at the Swiss National Supercomputing Centre (CSCS, Lugano, Switzerland) with 64 CPUs, taking less than 2 minutes in computing time. Effective strain was used as the output measurement for its ability to capture inhomogeneity in newly formed bone tissue. The dynamic character of the loading was not considered, in alignment with previous studies reporting that static simulations capture the main features and evolution of the mechanical environment^41^, while keeping computational complexity and time down.

To calculate the rescaled force suitable for the individual defect and healing progression, strain distributions were rescaled with a model intact vertebra with an 8N loading force used as the reference^30^. The 93rd percentile of strain resulting from the linear elastic FE model was used to rescale the loading force from 1% deformation to the intact reference strain level. It is accepted that the strain window most effective for bone modeling is between 800 and 2000 micro-strain^21^. Further, bone fails when 1 - 7% of the tissue units within the volume exceed 7000 micro strain^46^. To ensure that the strain distribution in the defect model satisfies both requirements, the 99th percentile of strain derived from five normal intact vertebrae of mice at similar age was used as a reference, and strain distribution in the defect vertebrae down-scaled by this factor (ratio=93^rd^ Defect/99^th^ Intact, Supplementary Fig. 1b). This rescaled force was individual to each mouse and adjusted after imaging (Supplementary Fig. S3), in what is termed real-time, as the bone is relatively unchanged during this period of scanning and computation.

### Animal model and loading

The first loading was applied two days after defect surgery, and three times per week thereafter. The defect vertebra was loaded via the pins in adjacent vertebrae using an in-house cyclic loading device, at 10Hz and 3000 cycles (5 minutes), similarly to previously reported^15^. The rescaled force calculated from the rtFE pipeline was used until the following weekly scan was completed, to balance the radiation exposure from imaging and the accuracy of the rescaled force. The rescaled force corresponded to the peak-to-peak cyclic loading, with 0.5N being the base position of the lower peak to avoid any inadvertent negative loads. Control mice were handled in a similar manner, and placed in the loading device immediately after their scan for 5 minutes without any loading applied. The control mice did not undergo the wait time associated with the rtFE computation.

### Analysis of bone defect healing

For evaluation, two regions of interest were defined from the baseline defect scan of each animal (Fig. 1a). The defect centre (DC) included the bony surface surrounding the defect (1 layer of voxels) as well as the space inside the defect. Region of interest determination was automated using a Hough transformation feature extraction technique by overfitting a cylinder to the medullary cavity and subsequently excluding any volume considered as existing bone within the created DC region. The defect periphery (DP) covered the remaining bone volume up to the start of the growth plates, as well as a dilated offset of 10 voxels to capture bone formation outside of the baseline cortical bone (Fig. 1a).

Bone volume fraction (BV/TV) was calculated in the DC (BV_DC_/TV_DC_) and in the DP (BV_DP_/TV_DP_) and normalized to their initial total volumes per region and per animal to calculate percentages^47^. Dynamic bone morphometric parameters were calculated in the DC and DP regions by registering binary micro-CT images acquired at consecutive weeks which were overlaid to compute regions of formation (F), quiescence (Q) and resorption (R) that could be analyzed morphometrically to yield bone formation rate (BFR) and bone resorption rate (BRR)^16,17^ in the DC (BFRDC and BRRDC) and in the DP (BFRDP and BRRDP), which is normalized to the respective initial volume, to calculate percentages per week, as shown previously^48^.

After registration, images were gauss-filtered (σ = 1.2, support 1) and thresholded to a binary image at 395mgHA/cm3 (corresponding to 4 GPa). Effective strain relevant to the (re)modelling events on the bone surface was estimated using FE analysis as per the rtFE solving pipeline. By combing surface (re)modelling events with corresponding surface effective strains, correlations and conditional probabilities could be investigated in the DC and DP, similarly to previously reported^15^.

Linear mixed-effects modelling was used for the statistical analysis (SPSS 24.0.0.0). Fixed-effects were allocated to: the time; and treatment. Random-effects were allocated to: the animal, to account for the natural differences in healing between different mice; and the animals’ specific defect volumes. Assumptions were tested by analyzing the residuals of the fitted model. Post-hoc tests with multiple pairwise comparisons were corrected with Bonferroni criteria. Data is reported as mean (±SD), unless otherwise stated. Reporting of statistics follows guidelines from the Publication Manual of the American Psychological Association (APA). A p-value of p < 0.05 was considered statistically significant unless reported otherwise.

### Data availability

All necessary data generated or analyzed during the present study are included in this published article and its Supplementary Information files (preprint available on bioRxiv 2020.09.13.295402; doi: https://doi.org/10.1101/2020.09.13.295402). Additional information related to this paper may be requested from the authors.

## Acknowledgements

The authors thank Nicholas Ohs, Duncan C. Tourolle for computational assistance, Ariane C. Scheuren and Charlotte Roth for animal assistance, and the Swiss National Supercomputing Centre (CSCS) for computational time. The authors thank Dr. Dr. Esther Wehrle for serving as deputy animal study director for this project providing advice on study design, surgery training, and veterinary support druing the animal experiments. AM is also grateful for the funding from the William Harvey International Translational Research Academy (WHRI-Academy, PCOFUND-GA-2013-608765), and the European Commission under the Marie Curie Co-funding of Regional, National and International Programmes (COFUND). The research leading to these results has received funding from the People’s Program (Marie Curie Actions) of the European Union’s Seventh Framework Programme (FP7/2007-2013) under REA grant agreement number 608765 This project also has received funding from the European Research Council (ERC) under the FP7/2007-2013, entitled Advanced MechAGE, ERC-2016-ADG-741883.

## Author Information

### Affiliation

Institute for Biomechanics, ETH Zurich, Zurich, Switzerland.

Angad Malhotra, Matthias Walle, Graeme R. Paul, Gisela A. Kuhn, Ralph Müller.

### Contributions

The study was designed by AM, GAK, and RM. The computational rtFE aspects were developed by AM, MW, and GRP, and performed remotely by MW. The animal experimental aspects were developed and performed by AM and GAK. Statistical and data analyses were performed by AM and MW. The manuscript was prepared by AM, and all authors reviewed and approved the final manuscript.

## Additional Information

Competing interests statement

The authors declare no competing interests.

