## Supplementary Information for "Application of subject-specific adaptive mechanical loading for bone healing in a mouse tail vertebral defect"

Institute for Biomechanics

ETH Zurich

Leopold-Ruzicka-Weg 4

8093 Zurich, Switzerland

### Supplementary Information

a

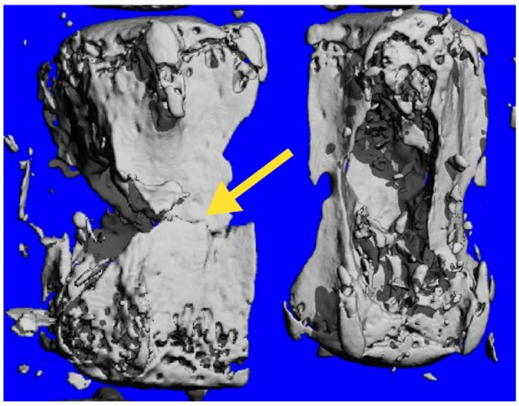

b

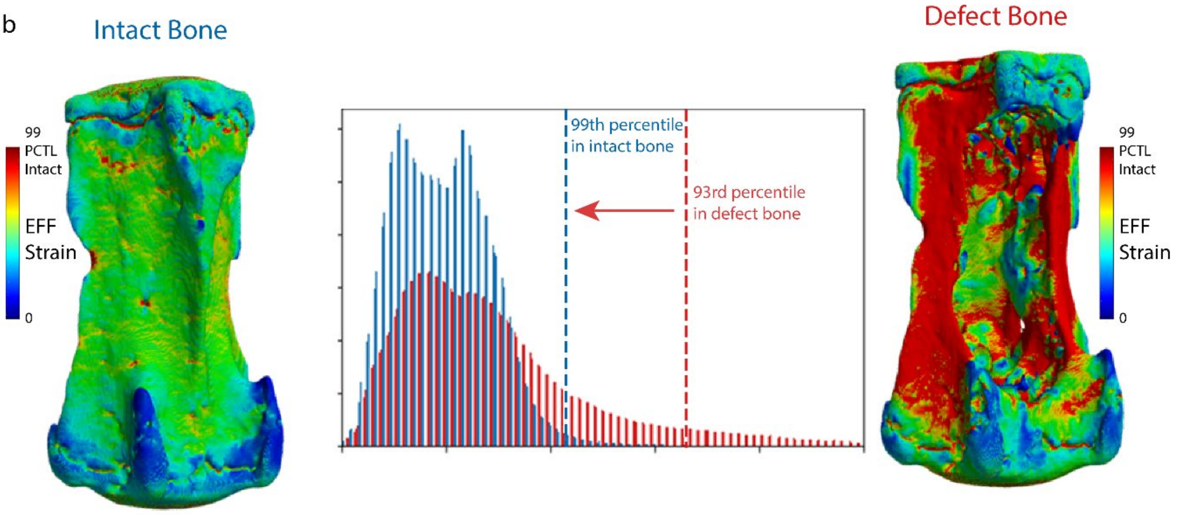


**Supplementary Fig. S1. (a)** Example of the risk of defect fracture. During pre-study testing of the surgery and the loading device in cadaver mouse vertebrae, at 15N the bone fractured around the defect (yellow arrow). This further justifies the need for rescaling of forces according to the defect severity and shape. Note: this was not part of the study and did not occur in a live animal. Due to this risk, and to factor in the initial defect and healing progression, loading regimes were downscaled. **(b)** Rescaling of loading conditions for a bone defect were considered from loading conditions shown to be effective for intact bone and downscaled to account for the increased effective strain around a bone defect.

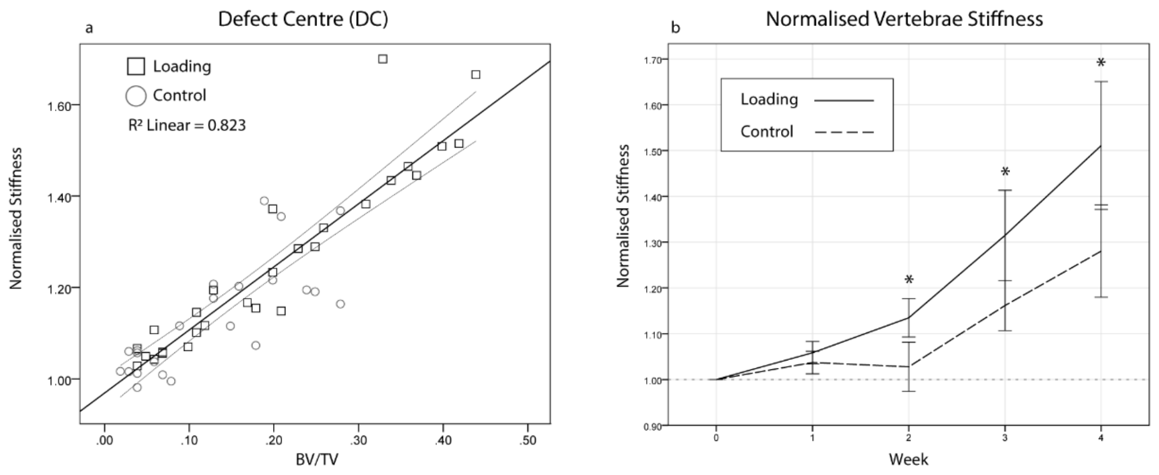


**Supplementary Fig. S2. (a)** Normalized stiffness was positively correlated to the defect centre (DC) bone volume fraction at week 4 (BV/TV), both in controls and in loading groups. **(b)** The vertebrae stiffness, assessed computationally, also significantly increased from week 2 onwards over controls.

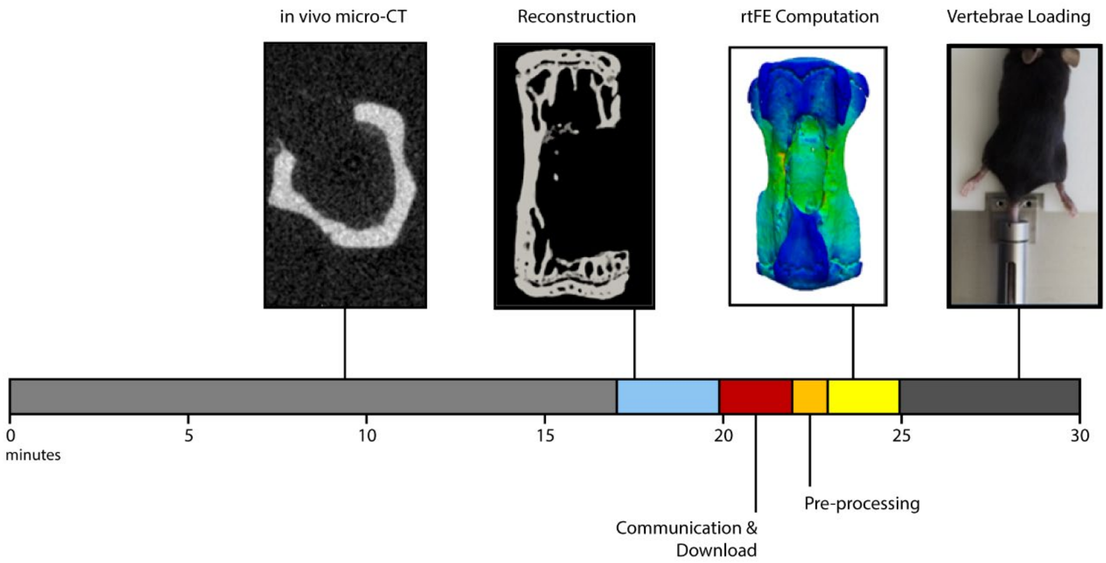


**Supplementary Fig. S3.** Timeline for rtFE workflow from when the mouse was anesthetized. The grey and dark grey regions were common to all animals. The light blue, red, orange, and yellow were specific to the rtFE methods, and completed remotely while the animal was kept under anesthesia and prepared for the loading device.
